# Adaptation to sub-optimal hosts is a driver of viral diversification in the ocean

**DOI:** 10.1101/261479

**Authors:** Hagay Enav, Shay Kirzner, Debbie Lindell, Yael Mandel-Gutfreund, Oded Béjà

## Abstract

Marine cyanophages are viruses that infect oceanic cyanobacteria, thus affecting global ecological processes. Cyanophages of the *Myoviridae* family are of great interest since they include generalist viruses capable of infection of a wide range of hosts including those from different cyanobacterial genera. While the influence of phages on host evolution has been studied previously, it is not known how the infection of distinct hosts influences the evolution of cyanophage populations. In marine systems this question is of special interest as the abundance of different *Synechococcus* and *Prochlorococcus* hosts constantly changes, temporally and spatially. Here, using an experimental evolution approach, we investigated the adaptation of multiple cyanophage populations to three distinct cyanobacterial hosts. We show that when infecting an “optimal” host, whose infection is the most efficient, phage populations accumulated only a few mutations. However, when infecting “sub-optimal” hosts, different, largely host-specific sets of mutations, spread in the phage populations, leading to rapid diversification into distinct subpopulations. The mutations included insertions, deletions, SNPs and codon adaptations. Most of the mutations were found in genes encoding for proteins responsible for host recognition, attachment and infection, regardless of their evolutionary conservation. Based on our results, we propose a model demonstrating how shifts in bacterial abundance, which lead to infection of “sub-optimal” hosts, act as a driver for rapid diversification of phage populations.

---

Cyanobacteria of the genera *Prochlorococcus* and *Synechococcus* are the most abundant photosynthetic prokaryotes in oceanic environments, contributing greatly to primary production on a global scale^1-3^. Viruses infecting cyanobacteria (cyanophages) are considered to be a major cause of cyanobacterial mortality^4-6^, therefore acting as a selective force on host populations^7,8^. Phages further promote cyanobacterial evolution, as they serve as agents for horizontal gene transfer^9^. Moreover, they are thought to play a role in biogeochemical cycling of dissolved and particulate organic matter^6,10^.

Cyanophages can be classified into one of three morphologically defined groups: *Podoviridae*, *Siphoviridae* and *Myoviridae*^11-13^. While cyanophages of the first two groups are mostly host-specific, many cyanophages of the T4-like *Myoviridae* family (cyanomyophages) are generalist parasites, capable of infection of multiple cyanobacterial hosts, even of different genera^11,13,14^. However, it is likely that different hosts are infected with varying efficiency by the same cyanophage, as was demonstrated previously for phages infecting *Flavobacterium* and enteric hosts^15,16^.

In natural habitats, the abundance of cyanobacterial hosts often changes over space and time^17-20^, as a result of seasonal environmental changes or killing off of the host due to phage infection^21^. Moreover, host populations can evolve resistance to infections by a specific phage strain, decreasing the number of the potential host strains for a given phage^7,22^. Such changes in the abundance of host strains that are infected most efficiently by a specific cyanophage can strongly influence the ability of the phage to reproduce. Thus, in order to maintain their reproduction, specific phage lineages would need to adapt to new hosts in their environment or else face the possibility of becoming extinct.

Previous studies that focused on the reciprocal co-evolution between hosts and phages have shown that this type of evolution results in rapid diversification^8,15,23^. A study that examined the one-sided evolution of a generalist phage, adapted to heat stress in two different enteric bacterial hosts, identified convergent evolution in phage populations^24^. However, the question of phage adaptation to optimal versus suboptimal hosts has yet to be addressed in environmentally relevant systems.

In this study, we investigated the adaptation of replicate cyanophage populations to different cyanobacterial hosts, in order to reflect rapid shifts in the abundance of hosts in natural environments. Using experimental evolution of cyanomyophage populations in three cyanobacterial hosts from two different genera, we investigated the extent to which the optimality of the host shapes phage evolution. In this context, optimality relates to the efficiency with which the phages infect their cyanobacterial hosts. Our results show, at the genomic and phenotypic levels, that infection of sub-optimal hosts is a major driver of viral evolution. This suggests that interactions with different host types is constantly shaping the structure of phage populations in natural environments, where host ecotypes coexist and form dynamic communities.

To examine how generalist cyanophages adapt to optimal and sub-optimal hosts we performed an evolutionary experiment whereby evolving phage populations were used to infect naïve cyanobacterial hosts. This was done for 15 rounds of serial infection which we estimate to have comprised thousands of viral genome replication events. A single isolate of the generalist S-TIM4 myovirus^7^ was used to infect three different cyanobacterial strains: *Prochlorococcus* sp. strain MIT9515, *Prochlorococcus* sp. strain MED4 and *Synechococcus* sp. strain WH8102 (referred to from now by genus and strain names). This was done with 4-5 replicate viral populations for each interaction with the different hosts. In order to avoid co-evolutionary dynamics and maintain the selective forces imposed by the host constant throughout the experiment, remaining cells were removed from phage lysates prior to initiation of the next round of infection (Figure 1).

**Figure 1.**
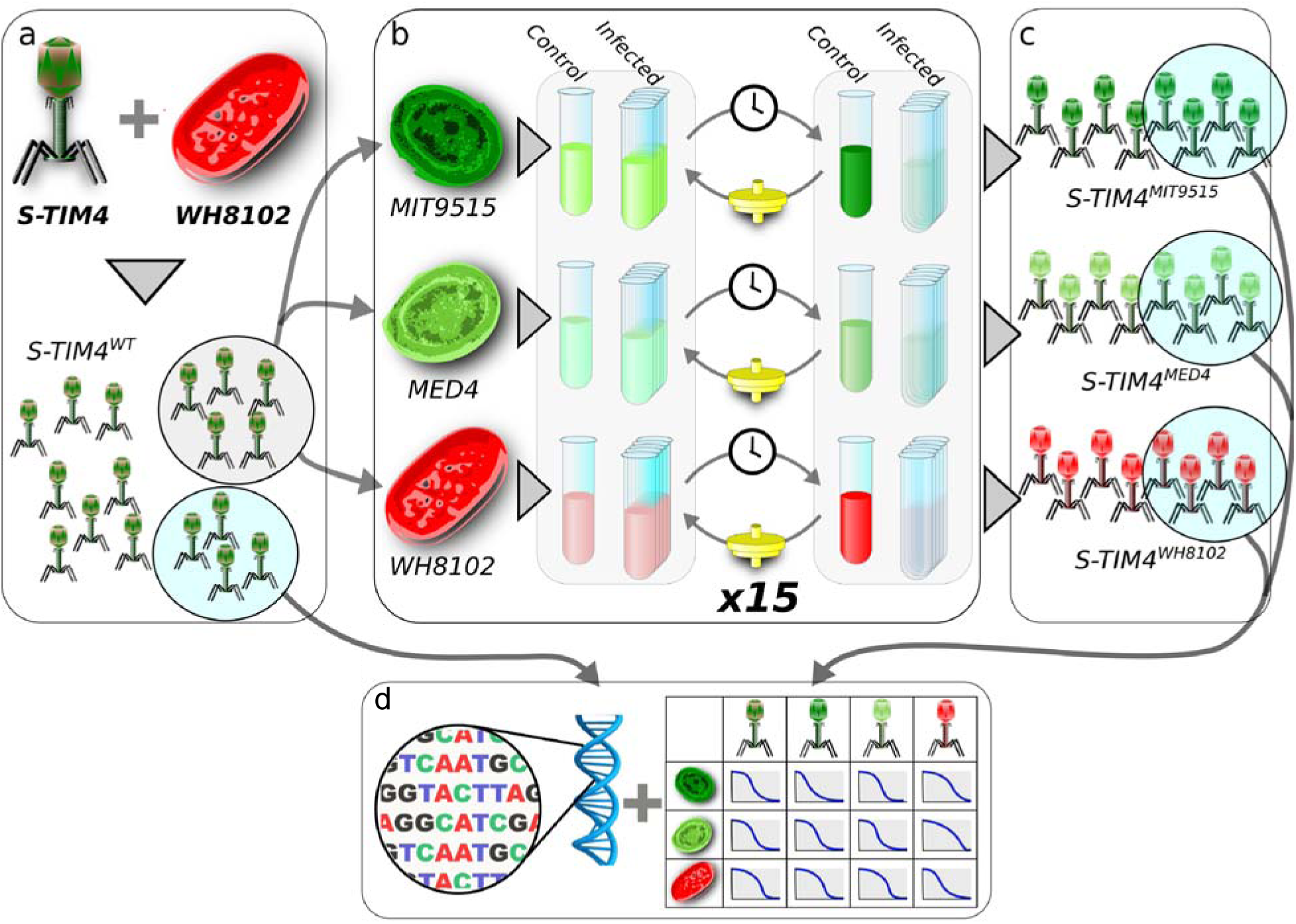
A schematic representation of the procedure for the experimental evolution of cyanophage S-TIM4 populations. (**a**) Generation of the ancestral population. A single viral plaque was isolated and replicated on the *Synechococcus* sp. strain WH8102 host, to yield a wild-type phage population (S-TIM4^WT^). (**b**) Adaptation to specific hosts. Aliquots of the wild-type population were evolved on 3 distinct cyanobacterial hosts: *Prochlorococcus* sp. strain MIT9515 (dark green), *Prochlorococcus* sp. strain MED4 (light green) and *Synechococcus* sp. strain WH8102 (red). Each infection interaction was repeated 5 times (Infected) with one uninfected control culture (Control) for each cyanobacterial strain. Following complete lysis of the infected cultures, phage lysates (< 0.22 μm) were transferred to infect the same naïve cyanobacterial hosts. This process was repeated for an additional 14 transfers. (**c**) Evolved populations. The adaptation phase resulted in populations adapted to MIT9515 (S-TIM4^MIT9515^), MED4 (S-TIM4^MED4^) and to WH8102 (S-TIM4^WH8102^). (**d**) Characterization of ancestral and evolved populations. Whole genome DNA sequencing of the entire population for the wild-type and each of the evolved populations was conducted (left). In parallel the infectivity of each population on each of the hosts was tested (right).

Following the evolutionary experiment, we performed whole population genome sequencing, using the average mutation frequencies from two sequencing libraries per evolved population. Additionally, to test for changes in the infection efficiencies of each phage population on each host, we infected the three hosts with each of the wild-type and evolved phage populations.

The S-TIM4 phage was isolated on *Synechococcus* WH8102^7^ and carries 235 genes in a 176 kbp genome. Infectivity tests of the unevolved S-TIM4 phage shows that it infects *Prochlorococcus* MIT9515 with the highest efficiency (Supplementary Table S1). Therefore we refer to this cyanobacterium as the “optimal” host. *Prochlorococcus* MED4 and *Synechococcus* WH8102 are infected with reduced efficiency, and are considered as “sub-optimal” hosts, with *Synechococcus* WH8102 being infected least efficiently.

## Mutations observed in S-TIM4 populations are a result of positive selection

Overall, we sequenced 14 S-TIM4 populations and identified 151 mutations in 86 different genomic positions (Table S2) that were localized in 20 protein-coding genes (Supplementary Table S3). Ninety percent of the detected mutations were Single Nucleotide Polymorphisms (SNPs), out of which, 89% were non-synonymous (Figure 2a). As the fraction of non-synonymous mutations within coding regions is expected to be ~75% in the absence of positive or negative selection^25^, the significantly larger fraction we observed of 89% suggests that the emergence of the identified mutations is the result of strong positive selection (Fisher test p-value = 3x10^−3^). An additional 12 synonymous SNPs modify viral codons, so that the mutated codons could form full Watson-Crick pairing with the tRNAs encoded in the host genome (Figure 2a, discussed below). This provides further evidence supporting that the mutations identified in this study are the result of positive selection.

**Figure 2.**
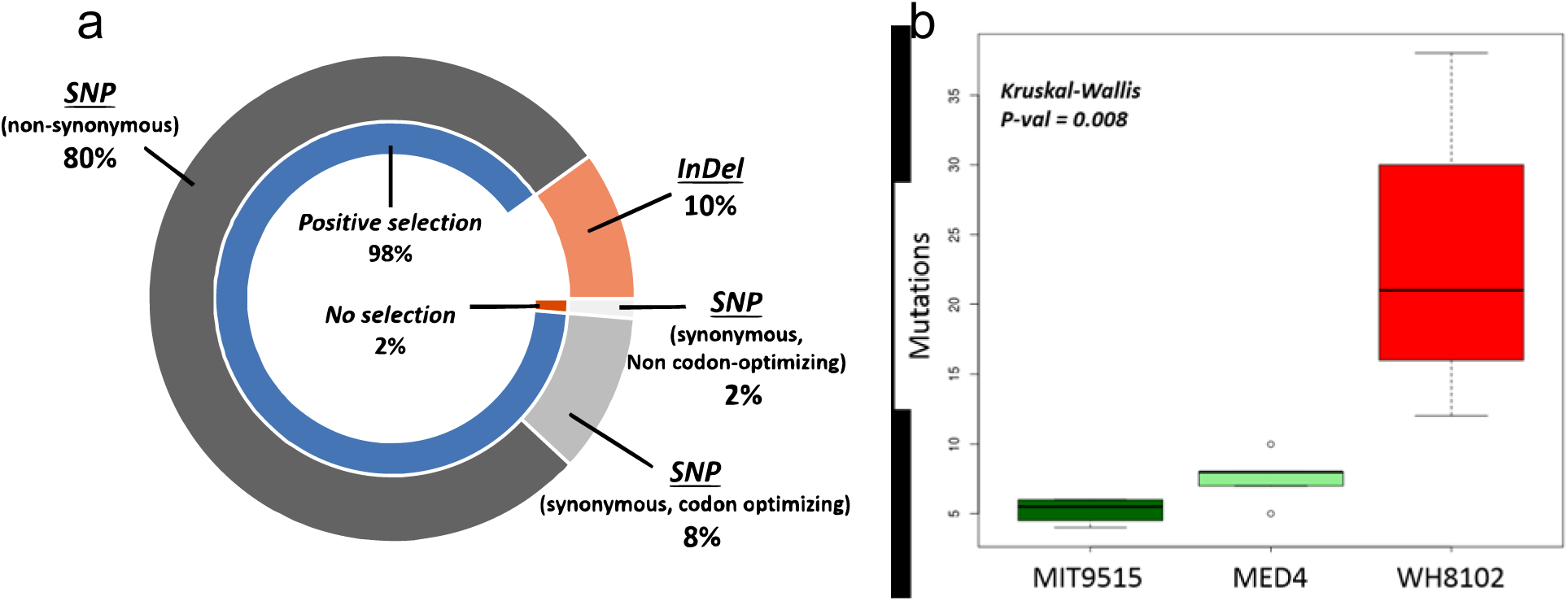
Mutations in evolved S-TIM4 populations are the result of strong positive selection. **(a**) Distribution of mutations type. Outer circle shows the distribution of mutations by type: insertions/deletions (orange), single nucleotide polymorphisms (SNPs) (grey scale: non-synonymous – dark grey, synonymous codon-optimizing for host tRNAs – medium grey, synonymous non-optimizing codon – light grey). Inner circle shows the distribution of SNPs by their potential to result in phenotypic change. (**b**) Distribution of the number of mutations accumulated in evolved populations, grouped according to the host they evolved in. Populations evolved in evolved in WH8102 (red) accumulated significantly more mutations, compared to populations evolved in either MIT9515 (dark green) or MED4 (light green) (Kruskal-Wallis p-value=0.008).

Examination of the number of mutations per population showed that S-TIM4 populations that evolved in the *Synechococcus* WH8102 host accumulated significantly more mutations, and in more genes, than populations evolved in the *Prochlorococcus* hosts (Figure 2b). The greater accumulation of mutations in these populations could be a result of cellular mechanisms of *Synechococcus* WH8102, making phages that were passaged through this host more prone to mutations. To examine this, we carried out an additional evolutionary experiment with cyanomyophage Syn19^26^ evolved on the same *Synechocccus* WH8102 host strain, for the same number of serial infection rounds. These populations accumulated 4-6 mutations per population. Therefore, we conclude that the greater accumulation of mutations in S-TIM4 populations passaged through WH8102 was not caused by the host cell per se, but was a result of a facilitated evolutionary process, resulting from the interaction between the phage and this sub-optimal host.

## Phage populations adapted to different hosts are phenotypically distinct

Next we sought to determine whether our phage populations that had evolved different genotypes had distinct phenotypes, as reflected by their infectivity profiles. The infectivity of each phage population was determined on each of the three hosts at two virus particle per cell ratios (VPC) of 0.1 and 3, as it has been demonstrated previously that the virus-host ratio influences infection efficiency^16^. We observed specialization of the evolved phage populations where populations evolved in *Synechococcus* WH8102 had improved infectivity of this host, compared to the wild-type population (Figure 3a & 3b). Furthermore, this came at a cost of a reduction in the infectivity of the two *Prochlorococcus* hosts. The reduced infection efficiency was so extreme in two of these populations that the ability to infect the MED4 host was completely lost (Figure 3a & 3b). In some cases, populations evolved on each of the *Prochlorococcus* hosts also evolved to infect that host better than the phages evolved on the other *Prochlorococcus* host. At times, this was at the cost of a reduction in the infectivity of the *Synechococcus* WH8102 host. Thus, we found that phage specialization was manifested as both improved infection of the host used for evolution and decreased infection of the other host types.

**Figure 3.**
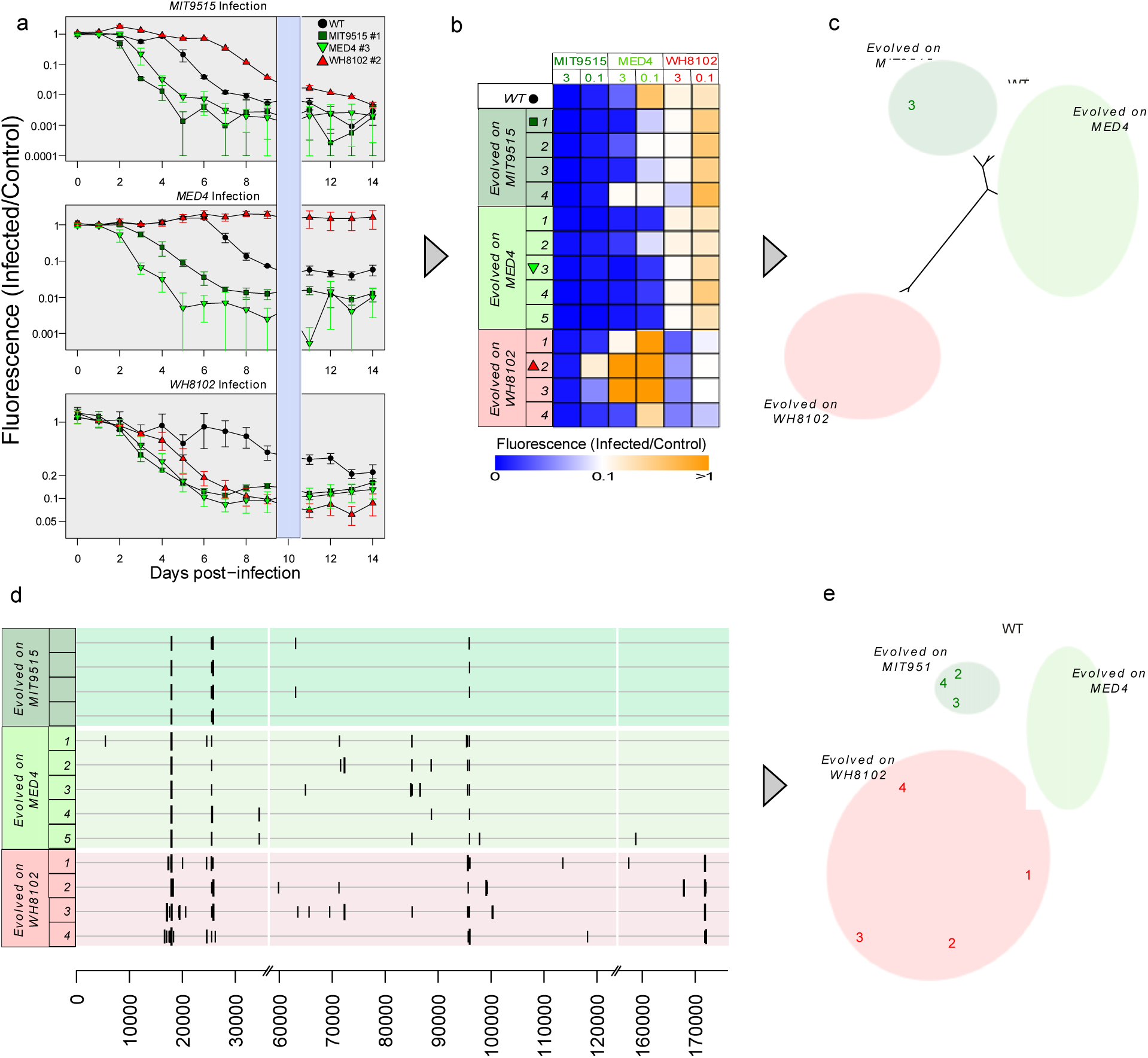
phage populations adapted to the same host show high within-group similarity. (**a**) Representative normalized growth curves of the three hosts when infected by four phage populations at a virus particle per cell ratio of 3. Phage populations shown are the wild-type (wt), evolved in the optimal host (MIT9515 #1), and evolved in sub-optimal hosts (MED4 #3, WH8102 #2). Y-axis shows the fluorescence levels of infected cultures divided by the fluorescence of the uninfected culture, i.e, normalized cell fluorescence. Blue shading shows normalized fluorescence on the 10^th^ day post infection, which was used for further analyses. Each curve is the average of 3 infection assays. (**b**) Normalized cell densities of all infected cyanobacterial cultures, 10 days post-infection. Lower values indicate more rapid declines in the infected host population, i.e. increased virulence. Rows correspond to viral populations, columns stand for infections of specific hosts at 0.1 and 3 viral particles/cell. Each box is the average of 3 infection assays. Symbols next to the population number correspond to the populations shown in panel a. (**c**) The phenotypic distance tree is based on the infection profile of each population (i.e., values of each row in panel b). Colors and numbering correspond to panel b. (**d**) Genomic maps show increased diversity in populations evolved in sub-optimal hosts. Positions along the genome are shown on the x axis; vertical lines correspond to specific populations. Horizontal lines correspond to identified mutations, and their length corresponds to the mutation frequency within the population. (**e**) Genotypic distance tree reveals lower distance in populations evolved in MIT9515. Tree was constructed based on pairwise genetic distances between phage populations. Colors and numbering correspond to panel b.

To further investigate the phenotypic difference between populations of phages that were passaged through different hosts, we computed a phenotypic distance tree based on the infectivity profiles 10 days post infection, at VPC of 0.1 and 3 (Figure 3c). Evolved phage populations clustered into three distinct groups, corresponding to the host they were evolved in. This clustering indicates that evolution of viral populations in the same bacterial host resulted in the most similar phenotypic profiles. Populations evolved in *Synechococcus* hosts clustered most differently to the wild-type phage population. Since *Synechococcus* was the least optimal host for the wild-type phage, these findings emphasize the increased phenotypic effect of adaptation to sub-optimal hosts.

## Phage populations evolved in sub-optimal hosts are genetically diverse

Next we sought a better understanding of how the phenotypic distances between phage populations that have evolved in different bacterial hosts are reflected in the mutational landscapes of the evolved populations. To do this, we created a genotypic profile for each phage population, containing the genomic positions and frequencies of all the mutations in the population (Figure 3d) and calculated the pair-wise genetic distances between each pair of phage populations and constructed a neighbor-joining tree (Figure 3e). Phage populations were clearly grouped according to the bacterial host they evolved on, in a similar manner to the structure of the phenotypic distance tree. However, the genetic distances varied between phage populations that evolved in the same bacterial host. Distances among populations that evolved in the optimal host, *Prochlorococcus* MIT9515, were the lowest, while for populations evolved in the least optimal host, *Synechococcus* WH8102, the distances were highest (Figure 3d & 3e, and Table S2).

Our combined findings suggest that strong purifying selection acted on mutations in S-TIM4 populations when evolving in the optimal MIT9515 hosts, with minimal genetic divergence occurring relative to the wild-type phage and the least diversity found among the different phage populations evolved in this host (Figure 3e). However, when S-TIM4 populations evolved in the non-optimal hosts, (*i.e*. *Synechococcus* WH8102 and *Prochlorococcus* MED4), the intensity of the purifying selection decreased, and positive selection resulted in the emergence of new diverse genotypes, with the greatest genetic divergence occurring during evolution on the least optimal host.

## Mutations in S-TIM4 are preferentially located within genes with a structural role

The majority of the 20 mutated S-TIM4 genes identified in this study are likely to be involved in building the viral particle. Based on homology to genes of the *Synechococcus* myophage Syn9^27^, many are predicted to be expressed during the last phase of viral gene expression (late-expressed genes) (Figure 4a and Supplementary Table S3), when most structural proteins are transcribed. Additionally, mass-spectrometry analysis of virus particles detected the proteins of 14 of the mutated genes (Supplementary Table S3), confirming that they have a structural role. Furthermore, all of the genes for these particle-associated proteins for which functions can be ascribed based on homology (8 genes) have structural functions^28^ (Supplementary Table S3). All but two of them have a common structural function of being associated with the tail fibers and baseplate, which are responsible for host recognition, attachment and infectivity^28,29^. Populations evolved in the *Synechococcus* host had additional mutations in genes encoding components of the tail tube and in capsid formation (Figure 4c). Of the six mutated genes that are not particle associated, two have putative functional predictions of involvement in the response to nutrient limitation. These are the 2OG^26^ and DUF680 (also referred to as PhCOG173) ^30^ genes.

**Figure 4.**
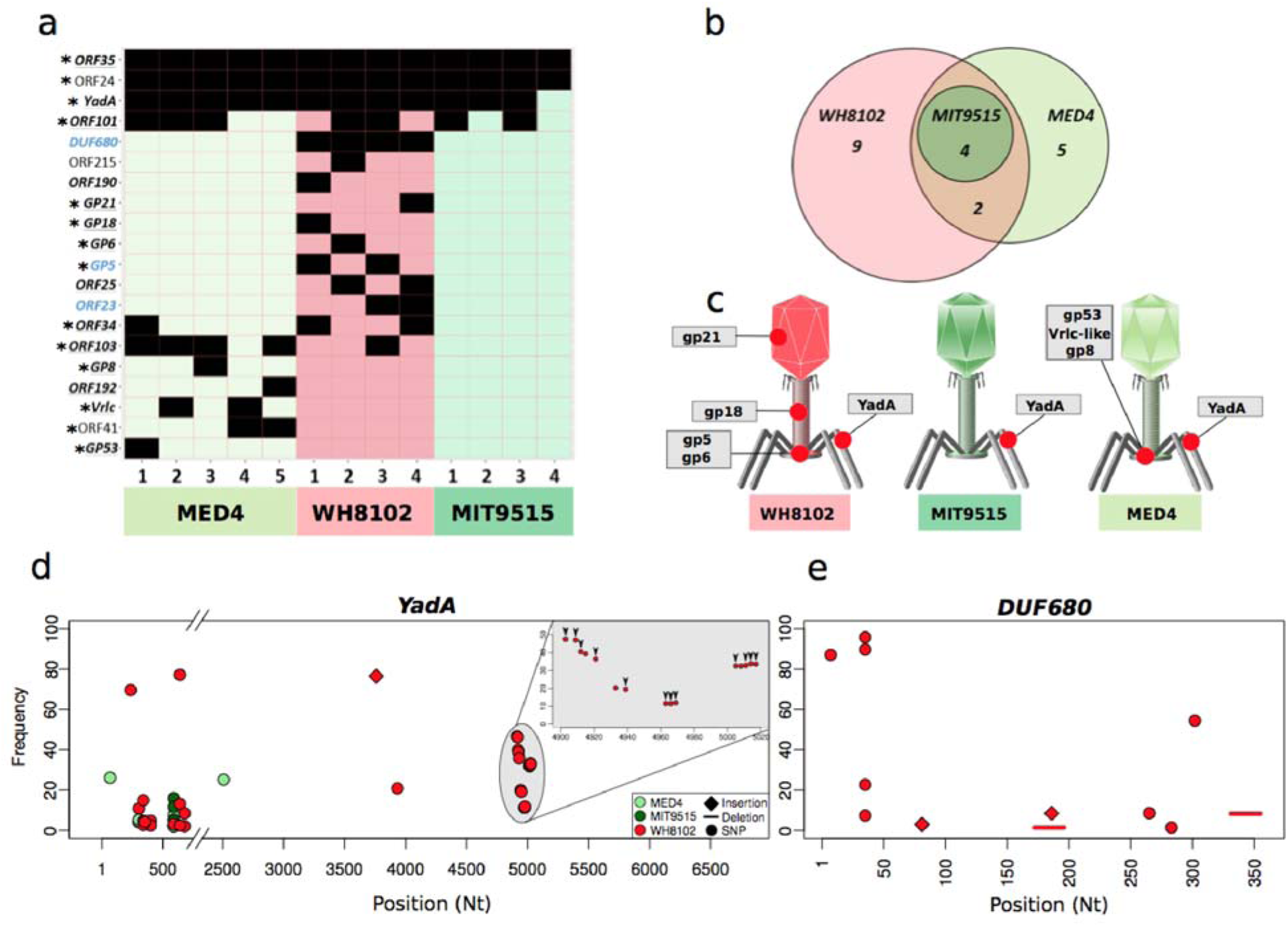
Discrete sets of structural genes are mutated in sub-optimal hosts. (**a**) **Mutated genes in each evolved population**. Each row represents one of the 20 mutated phage genes; columns represent evolved populations. Black cells stand for mutations identified in a particular gene. Genes in bold black type are predicted to be expressed during the late-phase of infection (and most have a structural role in the assembly of the virion particle). Genes in bold blue type are predicted to be middle-phase expressed genes. Genes marked with an asterisk encode proteins that were identified by mass spectrometry as part of the assembled viral particle. (**b**) Mutated genes in populations evolved in the optimal host are also mutated in other populations. Venn diagram showing the overlap of mutated genes in phage populations, grouped by the host they evolved in. (**c**) Location of mutated proteins in the virion particle. Each phage diagram represents populations evolved on a specific host: MIT9515, MED4 and WH8102. Protein names correspond to panel a. All proteins, excluding gp18 and gp21, are located in the tail fibers and baseplate, responsible for host recognition, attachment and infection. (**d**) Genetic map of the YadA gene (ORF108) shows increased diversity and specificity of mutations in populations evolved in suboptimal hosts. X axis shows the positions along the gene, mutation frequencies are on the Y axis. Grey inset plot is a zoom-in of a 120 base-pair long region which is enriched in 15 synonymous SNPs, all in the same population which evolved in WH8102 (#3). Black arrows are positioned over mutations which result in codon optimization to the host tRNA repertoire. (e) Genetic map of the DUF680-containing gene (ORF224) showing high diversity of mutations only in populations evolved in *Synechococcus* WH8102.

Mutations in four of the structural genes were common to the phage populations evolved on all three hosts (Figures 3d, 4). In fact, five of the same six mutations that were identified in the phage populations evolved in *Prochlorococcus* MIT9515 were also found in nearly all of the other phage populations (Figure 3d, Table S2). These findings suggest that these mutations emerged as an adaptation to a selective force imposed on all the cyanophage populations. This selective force could be a result of an intracellular component that is shared between hosts or possibly result from a feature of the extracellular environment (i.e. specific lab conditions). This hypothesis also explains why populations evolved on the optimal host often have improved infectivity of sub-optimal hosts, compared to the wild-type phage (Figure 3a, b).

The additional 16 mutated genes were found in phage populations that had evolved in either *Prochlorococcus* MED4 or *Synechococcus* WH8102, in sets of genes that were largely unique to each host (Figure 4a, 4b) with only two of the genes being common to populations evolved in both of these hosts. This indicates that genetic adaptation was, for the most part, distinctly tailored to each of the sub-optimal hosts in which the phage populations were evolved.

Next we provide two dramatically different examples of positive selection resulting in increased genetic diversity of phage populations during adaptation to sub-optimal hosts. The first is the YadA domain-containing structural gene that was mutated in all populations, and the second is the DUF680-containing non-structural gene (DUF, Domain Unknown function), that was mutated only in phages evolved in the *Synechococcus* WH8102 host.

The YadA domain-containing structural protein (2,235 aa long) has a typical membrane adhesion-like domain that likely has a structural role in the assembly of the viral tail-fibers that are involved in the initial phage attachment to the host cell^31,32^. This gene was mutated in a single position (Thr_580_=> Ala_580_) and at similar frequencies (5%-16%) in three of the four S-TIM4 populations passaged through *Prochlorococcus* MIT9515 as well as in populations evolved in the other two hosts. In contrast, mutations in the phage populations evolved in *Prochlorococcus* MED4 had 10 mutations in 4 distinct sites in the YadA gene. The divergence was even higher in phage populations evolved in the *Synechococcus* WH8102 host, where we found 33 mutations in 26 different sites, 31 of them different to those in the populations evolved in *Prochlorococcus* MED4 (Figure 4d). Strikingly, in one phage population passaged through *Synechococcus* WH8102 we identified 15 synonymous mutations that were restricted to a genomic region of 120 bp. Most of these mutations (12 out of 15) result in optimizing the codons to the host tRNA genes, potentially increasing the translation rate and accuracy of the YadA gene. These codon-optimizing SNP’s were mostly identified together on specific sequencing reads, meaning that specific viruses carry clusters of these mutations (Supplementary Figure S1). Previously it was suggested that clusters of non-optimal codons slow ribosome progression^33^, possibly changing protein structure and function as a result of different folding dynamics^34,35^. In a previous study we suggested that cyanomyophage genomes contain tRNA genes to allow improved translation of phage genes when infecting *Synechococcus* hosts^36^. The codon optimizing mutations we identified in this study represent a different mechanism to overcome the codon usage difference between cyanomyophages and their *Synechococcus* hosts.

The DUF680 protein (also referred to as PhCOG173) is common in cyanomyophages^37^ and was suggested to have a role in response to phosphate limitation^30^. This gene was mutated only in populations evolved in the *Synechococcus* host where it accumulated 13 different mutations, many of which are expected to interfere with its expression (*i.e*., frame-shifting indels, non-sense mutations and long deletions). Interestingly, in each population, the accumulative frequency of mutations in this gene is ~100%, suggesting that all phages in these populations carry a mutation in this gene. We speculate that the expression of the ancestral gene reduces the fitness of S-TIM4 when infecting *Synechococcus* WH8102 under the growth conditions used here. These two examples demonstrate starkly different means through which positive selection resulted in genetic diversification of phage populations, with the first likely improving the expression and functionality of the structural protein, while the second causes loss of function of a nutrient-response gene during adaptive evolution in nutrient-replete conditions.

## Adaptation to bacterial hosts occurs by mutations in conserved and variable genomic regions

Genes of cyanophages of the *Myoviridae* family can be divided into a conserved core-genome which is present in all phages of that group, and to a flexible, horizontally transferred genome, which is expected to allow adaptation to specific environmental conditions^37-39^. We asked if phage adaptation to its hosts is achieved by preferential accumulation of mutations in either the core or flexible genomes using the previous classification of cyanomyophage genes^37^. We found that 8 of our 20 mutated genes belong the core genome, i.e, they are included in all the cyanophage genomes of the *Myoviridae* group. Of the remaining 12 genes, 9 appear in some cyanophage genomes while only 3 genes have no homologs in other phages (Figure 5c).

**Figure 5.**
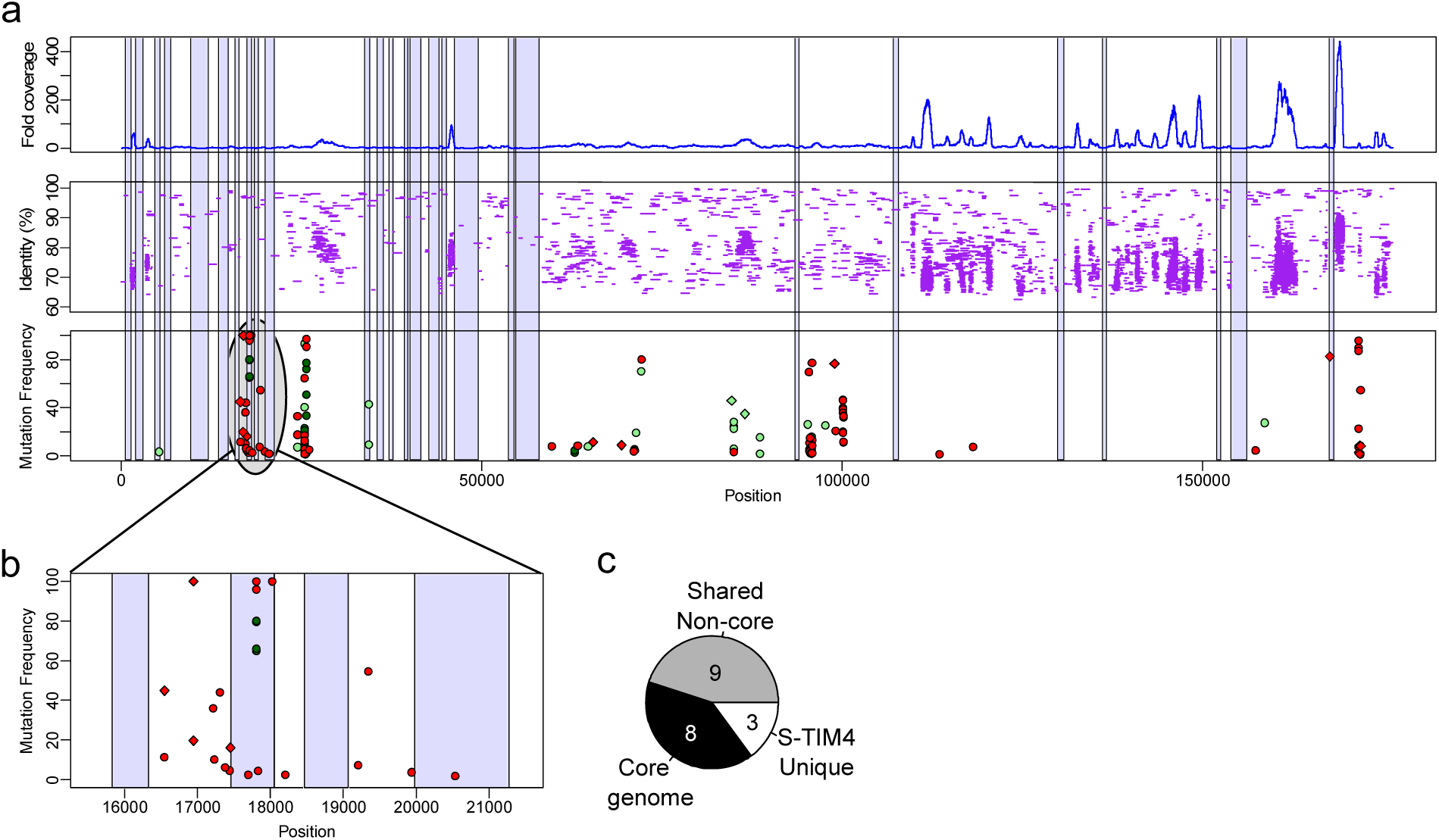
Phage adaptation to hosts occurs mainly through mutations in non-variable genomic regions and genes. (**a**) Mapping of metagenomic reads to the S-TIM4 genome reveals mutations in variable and conserved regions. The position along the genome is on the X-axis. The lower panel shows mutation frequency for combine phage populations per host along the genome. The middle panel shows the mapping of metagenomic reads to the S-TIM4 genome with the percent nucleotide identity of each read to the viral genome. The upper panel shows the fold-coverage of the metagenomic reads along the genome. Hypervariable genomic regions are shaded in blue (i.e. >500bp long, coverage lower than 20% of mean). (**b**) Zoomed in view of a mutation-rich genomic region (positions 15,500-21,500, open reading frames 23-27) of the S-TIM4 genome. (**c**) Adaptation of S-TIM4 to hosts occur by preferential mutation in genes found in other cyanomyophages. Genes are classified as unique in S-TIM4 (have no close homologs in other phages), belonging to the cyanomyophage core genome (present in all known cyanomyophage genomes), or shared non-core (appear in some cyanomyophage genomes).

Flexible genes often reside in hypervariable genomic regions^31^. Recently, it was suggested that genes responsible for host recognition and attachment reside within hypervariable genomic regions^31,39^. These regions are assumed to evolve rapidly to allow adaptation to new selective forces. This evolutionary adaptation occurs similarly in genomic islands described in bacterial genomes^40^, often resulting in bacterial resistance to infection by specific phages^7^. We therefore sought to determine if the mutations we identified in the S-TIM4 genome are preferentially located in hypervariable metagenomics regions using the Global Ocean Sampling (GOS) metagenome dataset^41^. Interestingly, only 14.5% of the S-TIM4 mutations are in hypervariable regions (hypergeometric P-value=0.728, Figure 5a, Supplementary Tables S2, S4). Therefore, we conclude that phage adaptation to specific hosts is not preferentially mediated through mutations in these hypervariable genomic regions. Overall, these data demonstrate that phage adaptation to a specific host is not gained by exclusive modification to either the core genome nor to the flexible genome.

## Model for the influence of host type on phage evolution

Based on our results, we propose a model for the influence of bacterial host type on the evolution of phage populations (Figure 6). According to our model, generalist cyanophages can infect a number of bacterial strains, with different degrees of efficiency, as was shown previously in other phage-host systems^15,16,42^. The bacterial strain infected with the highest efficiency is the “optimal” host, while other host strains are “sub-optimal”. When the availability of an “optimal” host is high, phage proliferation will occur mainly through infections of this host, as predicted by previous theoretical work^43^. As the phage is most adapted to such infections, the majority of the mutations that occur in the phage population result in lower infectivity (which likely directly reflects phage fitness), and their frequency is thus kept low as a result of purifying selection. Only a few mutations result in higher fitness when phages infect the optimal host strain, and the population converges closer to the maximal fitness point (Figure 6a). When availability of the optimal host declines either due to selective sweeps that result from environmental change, the acquisition of resistance by the optimal host, or the killing off of the host due to phage infection, then phage proliferation occurs by infection of sub-optimal hosts and the fitness of the phage population is expected to decrease. As a result, the population adapts to sub-optimal hosts: less mutations are eliminated by purifying selection and distinct sets of mutations are positively selected (Figure 6b, c). These mutations are mostly in genes responsible for host recognition, attachment and infection. Support for this part of our model comes from a recent study of phage adaptation to a new host in the mouse gut when the preferred host bacterium is absent^15^. The positive selection of genotypes carrying these mutation-sets results in rapid diversification. This would be the initial step in the separation of the phage population into distinct sub-populations.

**Figure 6.**
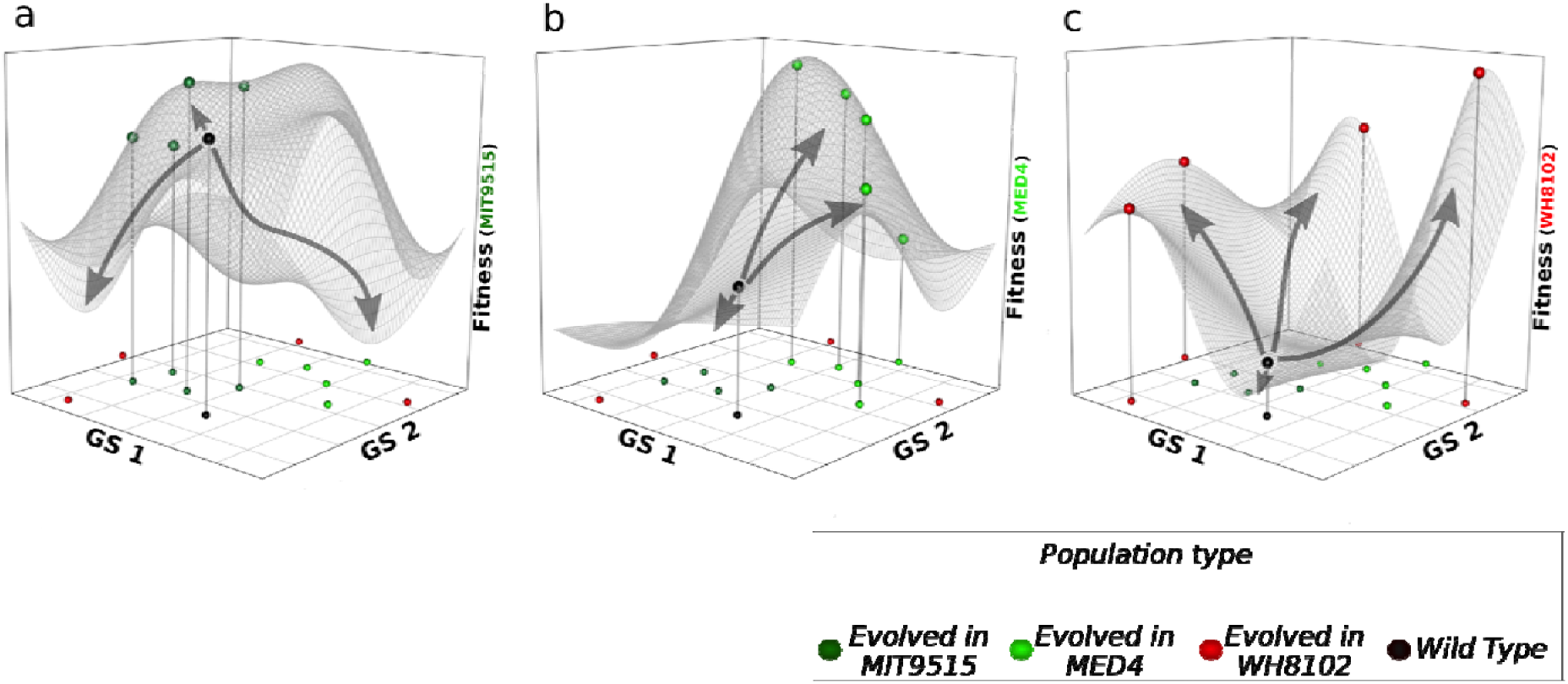
Model for the influence of different hosts on the evolution of viral populations. Each panel represents the infectivity landscape of the evolved and wild-type phage populations when infecting (a) the optimal host (i.e., infected most efficiently by the wild-type phage), *Prochlorococcus* MIT9515 (b) sub-optimal host *Prochlorococcus* MED4 (c) sub-optimal host *Synechococcus* WH8102. In all infectivity landscapes the GS1~GS2 plane (GS – Genomic Space) is identical and corresponds to pair-wise genetic distances between populations. The Z-axis shows the fitness of phage populations, based on the infectivity of each host (see methods). Predicted fitness surfaces (mesh over the genomic plane) were calculated based on the infectivity of all phage populations, when infecting each of the cyanobacterial hosts. Spheres located on the GS1~GS2 plane show the location of each phage population in the genotypic space. However, only the wild-type population and the populations that evolved in the host used to measure the infectivity appear as spheres on the fitness surface. When the optimal host is abundant, phage proliferation occurs mainly by infections of this host (panel a). The fitness of the wild-type population is high and most mutations result in lower fitness, therefore purified by negative selection (downwards arrows). Only a few mutations result in higher fitness and are positively selected (upwards arrow). Therefore, the evolved populations remain close to the wild-type population, both genotypically and phenotypically. Upon reduction in the abundance of this host, infections of “sub-optimal” hosts become frequent (panel b). When the fitness of the wild-type phages on this host is moderately lower, compared to the fitness in the optimal host (as happens in infections of *Prochlorococcus* MED4), phage evolution on these hosts results in a higher fraction of positively selected mutations, increased fitness of evolved populations (on this host) and higher genotypic diversification. Often, the only potential host is a bacterium, which is poorly infected by the wild-type phage (panel c). When the wild-type population evolves in this host high numbers of mutations are positively selected, leading to rapid divergence in phage populations, both phenotypically and genotypically. This divergence occurs mainly in genes responsible for host recognition and attachment, leading to separation into distinct phage populations, demonstrating how the type of bacterial host increases the diversity of phage populations.

A number of studies suggested that extinction of an abundant bacterial host and the proliferation of a rare host results in increases in the abundance of rare phage types (reviewed in reference^42^). We suggest, based on our findings, that changes in host availability not only change the abundance of different phages, but are also a key factor in the creation of the extensive degree of phage diversity observed in the environment.

## Acknowledgments

We would like to thank Sarit Avrani for providing cyanophage S-TIM4 and extensive help and Omer Nadel for preparing phage isolates for mass spectrometry analysis. This work was supported by the Louis and Lyra Richmond Memorial Chair in Life Sciences (to O.B.).

## Author contributions

H.E., Y.M.-G., D.L and O.B. designed the project. H.E. and S.K. performed laboratory experiments. H.E. performed bioinformatic analysis and wrote the manuscript with significant contributions from all authors. The authors declare no ing financial interests.

---

**Supplementary Figure S1.**
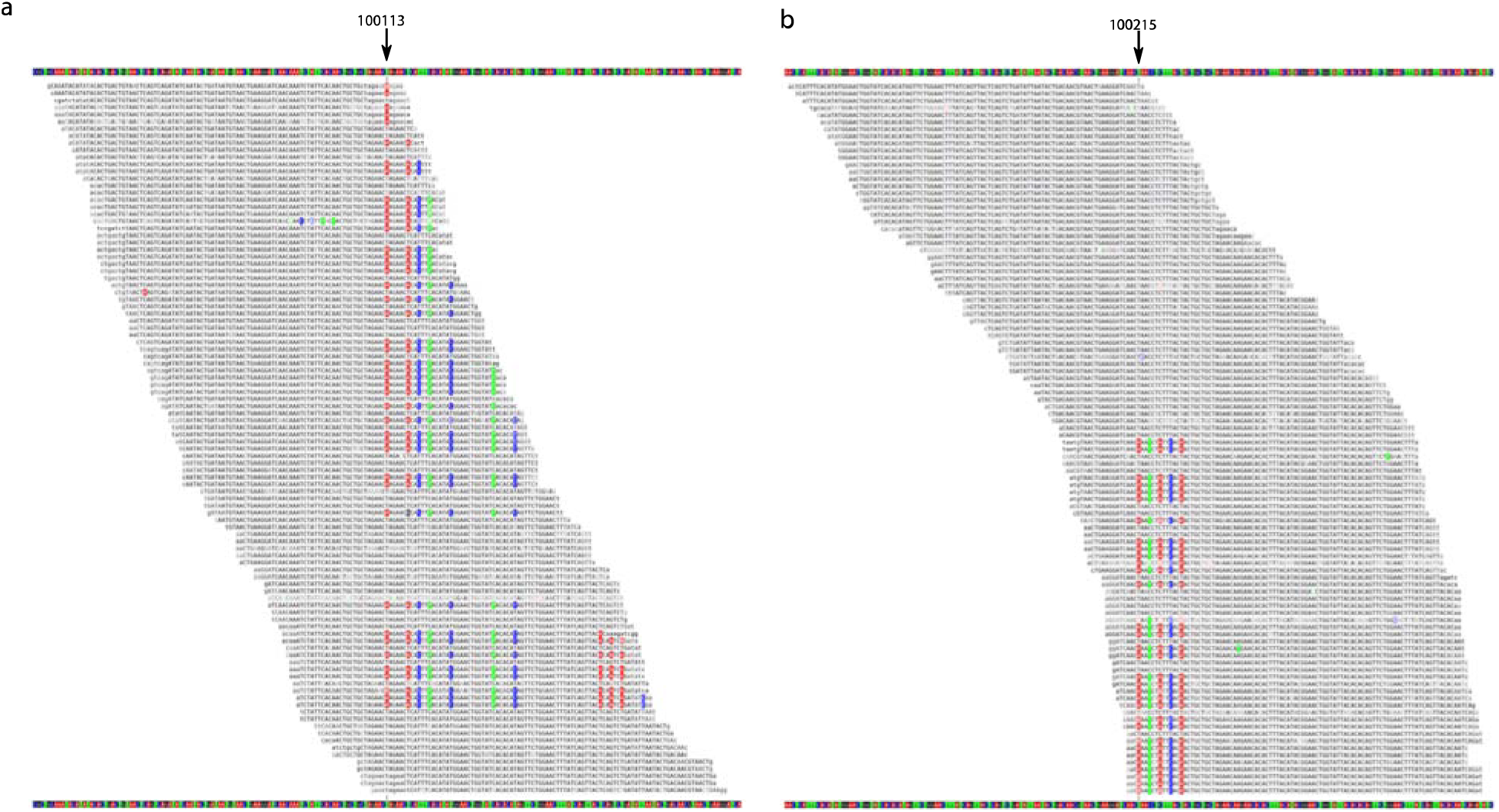
Codon optimizing mutations are clustered together in specific phage genomes. View of representative raw DNA reads, from population #3, evolved in WH8102 host. (**a**) Region flanking genomic position 100113. (**b**) Region surrounding position 100215. Highlighted bases represent mutations. Arrows on top show the genomic position of the central base in each genomic region view.

